# The Evolution of Protein Folds by Creative Destruction

**DOI:** 10.1101/2022.03.30.486258

**Authors:** Claudia Alvarez-Carreño, Rohan J Gupta, Anton S. Petrov, Loren Dean Williams

**Author notes:** Corresponding authors: Claudia Alvarez-Carreño, **Email:**, Anton S. Petrov, **Email:**, Loren Dean Williams, **Email:**.

## Abstract

Mechanisms by which new protein folds emerge and diverge pose central questions in biological sciences. Incremental mutation and step-wise adaptation explain relationships between topologically similar protein folds. However, the universe of folds is diverse and riotous, suggesting roles of more potent and creative forces. Sequence and structure similarity are observed between topologically distinct folds, indicating that proteins with distinct folds may share common ancestry.

We found evidence of common ancestry between three distinct β-barrel folds: OB, SH3 and cradle loop barrel (CLB). The data suggest a mechanism of fold evolution that interconverts SH3, OB and CLB. This mechanism, which we call creative destruction, can be generalized to explain other examples of fold evolution including circular permutation. In creative destruction, an open reading frame duplicates or otherwise merges with another to produce a fused polypeptide. A merger forces two ancestral domains into a new sequence and spatial context. The fused polypeptide can explore folding landscapes that are inaccessible to either of the independent ancestral domains. However, the folding landscapes of the fused polypeptide are not fully independent of those of the ancestral domains. Creative destruction is thus partially conservative in that a daughter fold would inherit some motifs from the ancestral folds. After a merger and refolding, adaptive processes such as mutation and loss of extraneous segments optimize the new daughter fold.

**Significance:** Mechanisms of emergence and early diversification of structured proteins present deep and difficult problems in evolutionary biology. Here we excavate the deepest evolutionary history, found within the translation machinery, which is an ancient molecular fossil and the birthplace of all proteins. We demonstrate common origins of some of the simplest, oldest and most common protein folds. Furthermore, the data suggest a mechanism, that we call creative destruction, that explains at molecular level how simple folds spawn more complex folds. In this mechanism, new folds emerge from old folds via gene duplication, expression, exploration of new folding landscapes and adaptation. Creative destruction explains the facile emergence of complex from simple architectures in a funneled exploration.

## Introduction

The simplest and most ancient protein folds are built from a small set of supersecondary structures (1). Folds with more complex architectures emerged from a funneled exploration; there is insufficient time and resources in the universe to find novel folds by random searching of sequence space (2).

Around a thousand folds form the basis of hundreds of thousands of proteins (3), generating the vast universe of function (4-6). A fold is a well-defined and specific layout of backbone topology and secondary structural elements (7). At the base of the protein structure hierarchy, polypeptide chains form intramolecular hydrogen bonds within α-helices, β-sheets and loops (8, 9). At the next level of the hierarchy, secondary structural elements combine to form supersecondary structural elements such as β-α-β, β-hairpin or helix-turn-helix (10-13). At even higher levels of the structure hierarchy, secondary and supersecondary structural elements assemble with each other to from compact globular folds (4, 14, 15).

Fold origins and evolution pose central questions in biological sciences. How did ancient folds arise? What evolutionary mechanisms led to the diverse set of protein folds in contemporary biological systems? Modest divergence from an ancestral fold can take place by incremental adaptive mutations that convert one type of secondary element to another (16). However small stepwise adaptive steps cannot explain large-scale fold rearrangements (17, 18).

Here, we describe a general mechanism of fold evolution, in analogy with Schumpeter’s “perennial gale of creative destruction” in economics (19). On the level of molecules, creative destruction provides a general and highly accessible pathway for diversification of structure and function. Creative destruction may account for the observed diversity of protein folds and affords experimental approaches to exploration of new fold space.

In creative destruction of protein folds, an open reading frame duplicates or otherwise merges with another (20) to produce a fused polypeptide (Figure 1a-b). A merger forces two ancestral domains into a new sequence and spatial contexts, with the possibly of physical impingement. A merger thus has the potential to destabilize or destroy the folds of the ancestral gene products. At the same time, the fused polypeptide might collapse to a new fold (Figure 1c). The fused polypeptide can explore folding landscapes that are inaccessible to either of the independent ancestral sequences. However, the folding landscapes of the fused polypeptide are not fully independent of those of the ancestral domains. Creative destruction is thus partially conservative in that a daughter fold would inherit some or all motifs from the ancestral folds and would also contain new elements (Figure 1d). After gene merger, polypeptide expression, and exploration of new folding landscapes, adaptive processes such as mutation and loss of extraneous segments optimize the new daughter fold.

**Figure 1.**
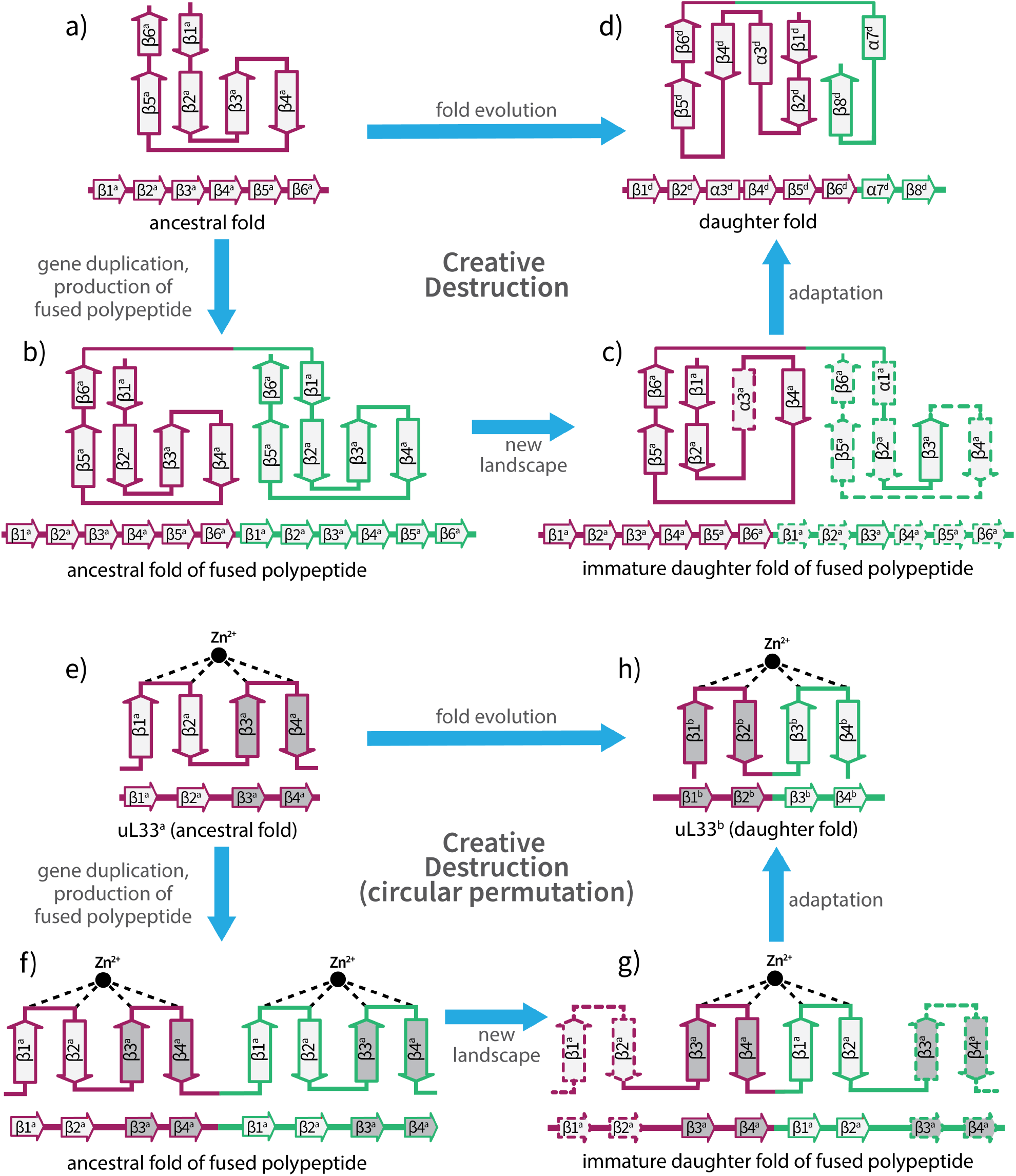
Creative destruction. Top: General mechanism of creative destruction; an ancestral fold is destroyed, and a daughter fold is created. This figure shows, on a topological level, (a) an ancestral fold, (b) the ancestral folds of the fused polypeptide, (c) the immature daughter fold of the fused polypeptide in which parts of the ancestral folds and some secondary elements have been destroyed and an immature daughter fold has been created, and (d) the mature daughter fold, which has inherited some but not all supersecondary elements of the ancestors. Bottom: The special case in which creative destruction causes circular permutation of ribosomal protein uL33. (e) an ancestral uL33 fold, (f) the ancestral folds of two fused uL33 polypeptides, (g) the immature circularly permuted daughter fold of uL33, and (h) mature circularly permuted uL33. A duplication of β1^a^β2^a^β3^a^β4^a^ gives the fused polypeptide β1^a^β2^a^**β3**^**a**^**β4**^**a**^ -- **β1**^**a**^**β2**^**a**^β3^a^β4^a^, (where -- is a linker). The circles represent zinc ions. Strands are selectively shaded to facilitate tracking them through the creative destruction process. The fused polypeptide folds in a new landscape and resolves by adaption (∼ indicates cross-fold sequence similarity). The dashed secondary elements are unstable in the immature daughter folds and are lost in the mature daughter fold. The ancestral folds of the fused polypeptide are included in the schematic to illustrate destruction of the ancestral folds and inherence of some ancestral secondary motifs.

In creative destruction, well-characterized and frequent genetic processes (20) provide access to partially conservative new folding landscapes without implausible combinatorial shuffling of supersecondary structures. For simplicity, the model here focuses on gene duplication. However, truncation, or deletion, or incorporation of exogenous coding or non-coding sequences might initiate creative destruction.

Circular permutation is a special case of creative destruction and is a common and explanatory example of fold evolution (Figure 1e-h). Inspection of structures of initial ancestral and final daughter folds might suggest that circular permutation is accomplished simply by rearrangement of linkages between secondary structural elements. In reality, the mechanism involves gene duplication and expression of a fused polypeptide that fails to fold to the ancestral domains but does fold in a new landscape with partial conservation of secondary elements. Here we illustrate how zinc-binding ribosomal protein uL33 (21) is circularly permutated by creative destruction. The mechanism entails duplication and in-frame fusion of the uL33 gene (16, 22), fold destruction, fold creation (in a new landscape), and adaptation (Figure 1g). The secondary elements of the two ancestors are semi-conserved in the daughter fold (half of them are conserved and the other half are lost, Figure 1h).

We provide sequence and structural support for widespread occurrence of creative destruction during fold evolution. Our focus is on some of the oldest, simplest, and best-preserved proteins in biology. Vestiges of creative destruction are seen by comparisons of three ancient β-barrel folds (Figure 2): Oligonucleotide/Oligosaccharide-Binding, OB (23); Scr kinase family Homology 3 (SH3); and cradle loop barrel, CLB (24). Proteins with OB, SH3 and CLB folds are found in central metabolic processes and throughout the translation system, including in ribosomal proteins, translation factors, and aminoacyl tRNA synthetases. We use CLB to refer to the Alanine Racemace C topology of the cradle loop barrel fold (25). Our results reveal creative destruction is a mechanism of circular permutation, and explain common ancestry of SH3, OB, and CLB fold proteins.

**Figure 2.**
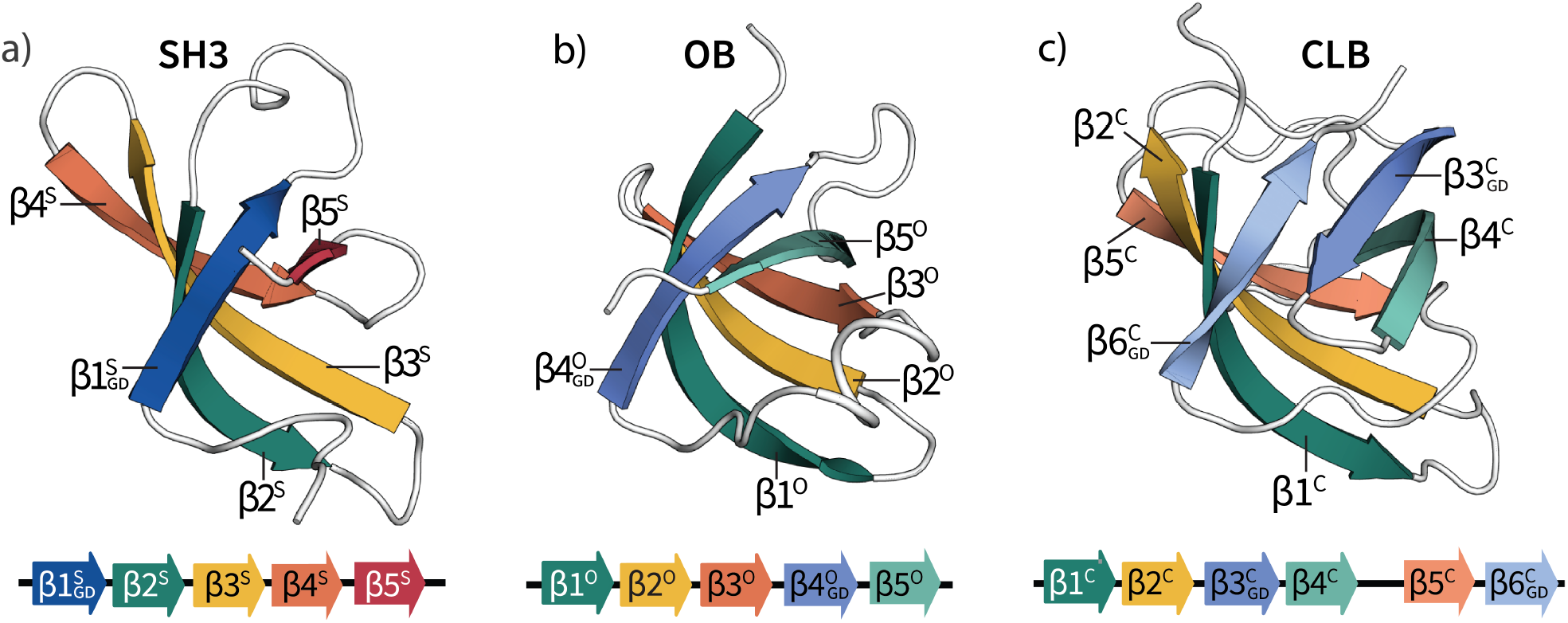
Representative structures of OB, SH3 and CLB folds. (a) Structure of an SH3 fold (PDB: 1NZ9, chain A). (b) Structure of an OB fold (PDB: 2OQK, chain A). (c) Structure of a CLB fold (PDB: 4B43, chain A). OB and SH3 are five-stranded β-barrels, CLB is a six-stranded β-barrel. GD: GD-box motif. The color scheme suggests common ancestry.

## Results

Three-dimensional structures of biopolymers are generally more conserved than sequences (26, 27). Exceptions to this pattern are called cross-fold sequence similarities, which are sequences that are conserved between proteins with different three-dimensional structures. Cross-fold sequence similarities indicate shared evolutionary history between distinct folds and are recognized as evidence of fold evolution (11, 16, 28, 29). In some cases, distinct folds reveal multiple regions of sequence similarity that are permuted relative to each other.

We have detected homologous relationships within and between SH3, OB, and CLB folds. To improve the sensitivity of the sequence-search method, we used protein sequence profiles instead of single-sequence searches, following the workflow in Figure 3 (see Methods). Protein sequence profiles capture the distribution of conserved and non-conserved positions across a multiple sequence alignment and improve the sensitivity of homology detection (30). The HHalign score (30) indicates the estimated probability of homology between a query and a template profile.

**Figure 3.**
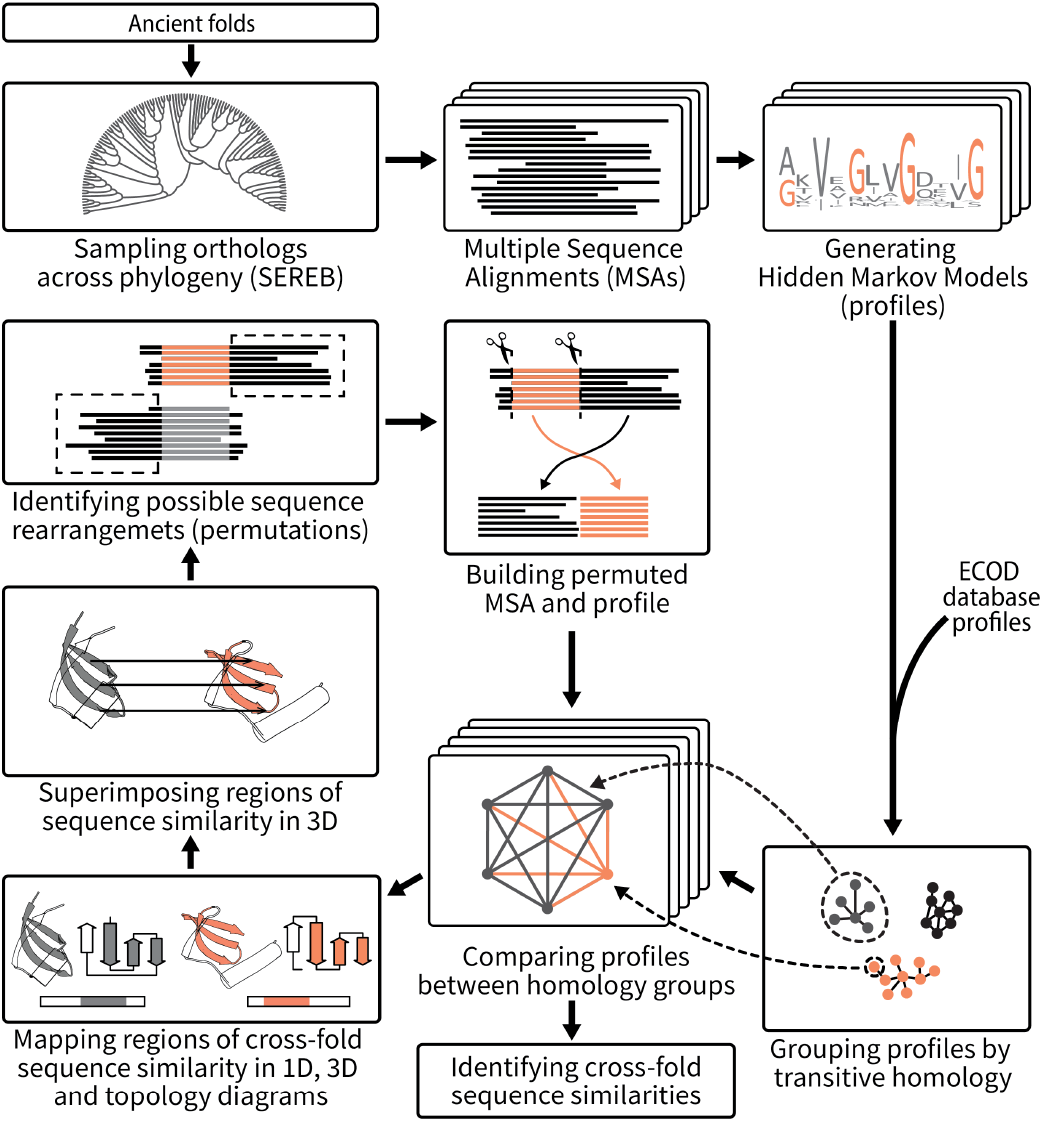
Workflow for identifying evolutionary relationships between folds. Orthologous protein sequences for a given fold within the SEREB database were aligned. Multiple sequence alignments (MSAs) were used to create profile Hidden Markov Models for each fold. Profile-profile comparisons between folds in the ECOD database revealed regions of sequence similarity. The method revealed transitive homologous relationships. The method also revealed cross-fold similarities covering distinct and, in some cases, overlapping regions between distinct folds.

We consider HHalign scores higher than 50% to be indicative of shared common ancestry within a given fold. In several instances, two sequence profiles are seen to be homologous to each other by way of homology to a common third sequence (31). This phenomenon, called transitivity (32), has facilitated detection and characterization of signals of creative destruction.

We consider HHalign scores higher than 60% to be indicative of shared common ancestry between regions of different folds (cross-fold sequence similarity). The patterns of cross-fold sequence similarities between distinct representatives of SH3, OB and CLB folds are consistent with, and provide strong support for, creative destruction as a mechanism of fold evolution.

In addition to sequence similarity, we analyze structural similarity between SH3, OB, and CLB, including, β-sheets, β-hairpins and GD-box motifs (33, 34). A GD-box is a motif that contains a β-strand connected to a loop by a β-turn, and portions of a second non-contiguous β-strand (33). A GD-box is characterized in part by the amino acid sequence ΨxΨxxGρxΨxΨ, where G is glycine, Ψ is aliphatic, ρ is polar and x is anything.

To understand and explain creative destruction, we use a reduced representation of protein structure: 1) α refers to an α-helix, β refers to a β-strand, and L refers to a loop; 2) a number following an α or β indicates the relative position in the sequence, N to C; 3) secondary structural elements are notated by fold as α° or β° or L° (indicates OB fold), α^S^ or β^S^ or L^S^ (indicates SH3), or α^C^ or β^C^ or L^C^ (indicates CLB); 4) L is only specified for loops that display cross-fold sequence similarity to an α-helix or a β-strand; 5) β_GD_ indicates a GD-box; 6) -- refers to a linker between two domains; and 6) ∼ indicates cross-fold sequence similarity between secondary elements. For example, β1° ∼ β3^S^ indicates the first β-strand of an OB fold protein has sequence similarity with the third β-strand of an SH3 fold protein.

### SH3 and OB Folds

OB and SH3 folds are both five-stranded β-barrels with two antiparallel β-sheets (23). Although the topology of SH3 and OB folds differs, β-strands and some of the connective elements overlay in three-dimensions (34-36). SH3 and OB folds share one region of cross-fold sequence similarity corresponding to β2^S^β3^S^β4^S^ ∼ β1°β2°β3° (Figure 4e) and a second region of cross-fold sequence similarity corresponds to β1 ^S^ β2_GD_^S^ ∼ β4_GD_° β5° (Figure 4f).

**Figure 4.**
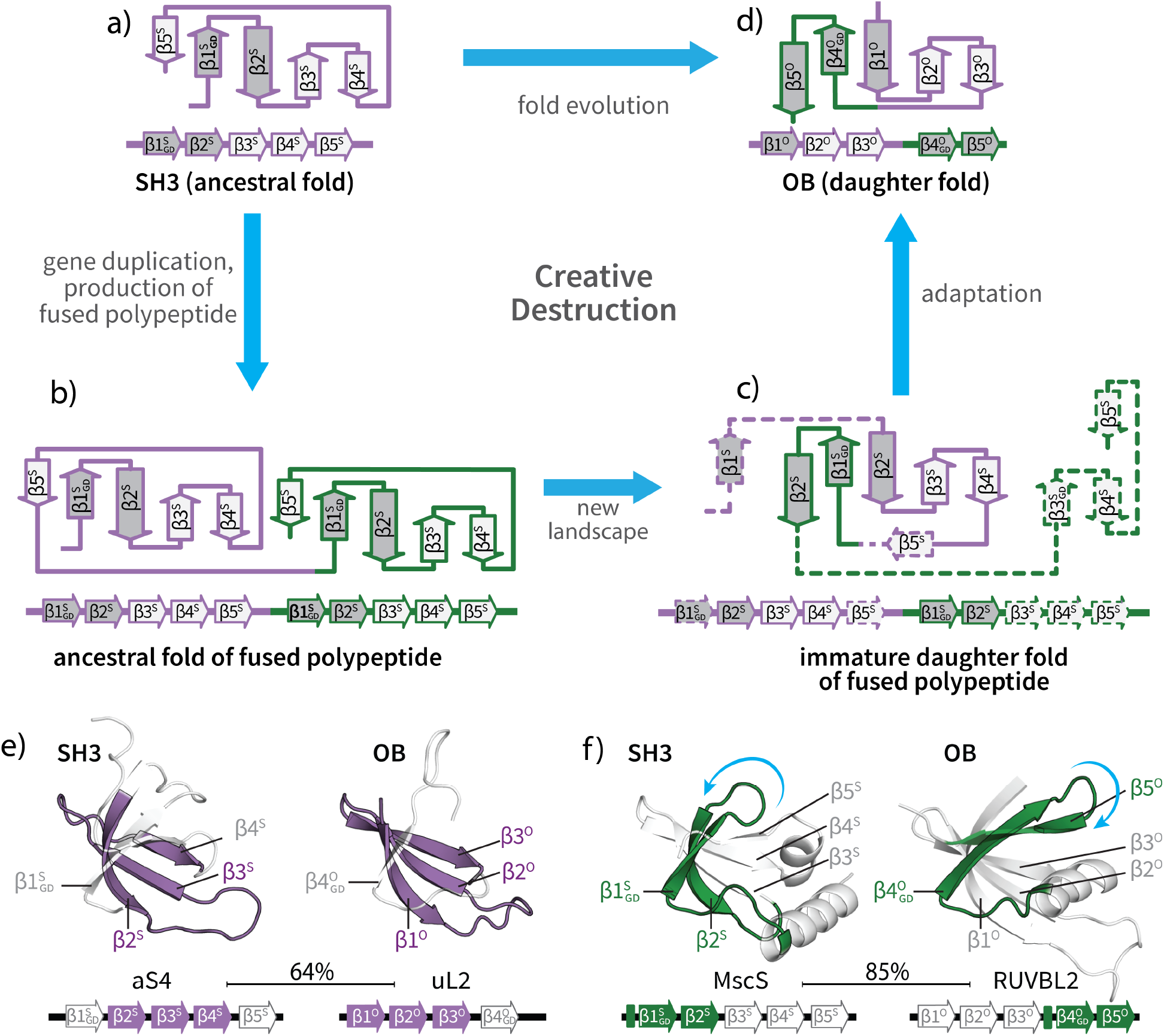
Conversion from SH3 Fold to OB Fold by creative destruction. This figure shows, on a topological level, (a) the ancestral fold of the native protein, (b) the ancestral folds of the fused polypeptide, (c) the immature daughter fold in the fused polypeptide, and (d) mature daughter fold. A duplication of SH3 gives the fused polypeptide SH3-SH3. The fused polypeptide refolds and resolves by adaption, yielding an OB fold. GD: GD-box motif. The ancestral folds of the fused polypeptide are included in the schematic to illustrate destruction of the ancestral folds and inherence of some ancestral secondary motifs. The fused polypeptide in reality would bypass the ancestral folds, and collapse directly to the immature daughter fold. β2^S^ ∼ β1°, β2° ∼ β3^S^, β4^S^ ∼ β3°, and β2^S^ ∼ β5° are shaded in dark gray allow tracking of β-strand positions during the transition from ancestral fold to daughter fold. (e) Cross-fold sequence similarity between aS4 and uL2 corresponds to antiparallel β-strands β2^S^β3^S^β4^S^ and β1°β2°β3°. (f) Cross-fold similarity between MscS and RUVBL2 corresponds to β1_GD_^S^ and β4_GD_° Mapping of sequence similarity between SH3 and OB: A bar between two folds indicates cross-fold sequence similarity, as given by HHalign scores. Regions that yield no sequence similarity are shown in white. Secondary structural elements that share sequence similarity are indicated by the same color in both members of the pair. Curved arrows follow secondary structural elements that are display differences in conformation between folds.

The patterns of cross-fold sequence similarity, topology, motif conservation and 3D structure between SH3 and OB support creative destruction (Figure 4a-d). A duplication of β1_GD_^1^β2^S^β3^S^β4^S^β5^S^ yields the fused polypeptide β1_GD_^S^**β2**^**S**^**β3**^**S**^**β4**^**S**^β5^S^ -- **β1**_GD_^**S**^**β2**^**S**^β3^S^β4^S^β5^S^. In our schematic, the fused polypeptide initially collapses to the ancestral folds, and then converts to an immature daughter fold. The collapse to the ancestral folds is shown for illustrative purposes only; the fused polypeptide actually collapses directly to the immature daughter fold, which resolves by adaption. The final product contains bold secondary elements **(β2**^**S**^**β3**^**S**^**β4**^**S**^**β1**_**GD**_^**S**^**β2**^**S**^), which are renumbered (N to C) to give β1°β2°β3°β4_GD_°β5°. Creative destruction is supported by cross-fold sequence similarities β1° ∼ β2^S^, β2° ∼ β3^S^, β3° ∼ β4^S^, β4_GD_° ∼ β1_GD_^S^, β5° ∼ β2^S^. This pattern of cross-fold sequence similarity is consistent with formation of an OB fold by duplication of an SH3 gene, polypeptide expression, collapse to an immature daughter fold, and adaption, to yield a SH3 fold (Figure 4a-d).

The simpler topology of SH3 compared to OB suggests that SH3 is the ancestor of OB. However, the opposite polarity, where a duplicated, refolded and adapted OB results in the emergence of the SH3 fold is also consistent with the data. We previously described how gene fusion, refolding and adaptation provide a simple mechanism of conversion between OB and SH3 folds (34). Below we provide support for each step of creative destruction using specific examples.

### Ribosomal proteins uL2 and aS4

Cross-fold sequence similarity is seen between ribosomal protein uL2 (OB) and ribosomal protein aS4 (SH3). These two proteins share one region of sequence (HHalign score 64%) and structure similarity of the antiparallel β-sheet (β2^S^β3^S^β4^S^ ∼ β1°β2°β3°, Figure 4e and Supplementary File 1, Table S4). Sequence similarity between uL2 (OB) and ribosomal protein aS4 (SH3) is not apparent in the antiparallel β-sheet that contains the N- and C-terminal strands.

### RUVBL2 and MscS

Cross-fold sequence similarity is seen in RuvB-like AAA ATPase 2 (RUVBL2, OB) and the mechanosensitive channel of small conductance (MscS, SH3) (Figure 4f and Supplementary File 1, Table S4). The region of sequence similarity is β1_GD_^S^β2^S^ ∼ β4_GD_°β5° (HHalign score 85%).

As predicted by creative destruction conversion of SH3 to OB requires refolding. β1_GD_^S^β2^S^ and β4_GD_°β5° are contained within different antiparallel β-sheets. β5° is on the edge of β°β°β° while β2^S^ (similar in sequence to β5°) is on the interior of β^S^β^S^β^S^. The relative position of β5° is the same as β5^S^ within the linear sequence, but not within the three-dimensional structure.

### SH3/OB summary

Homology relationships between OB folds and SH3 folds can be generalized. We have shown that uL2 and aS4 (β2^S^β3^S^β4^S^ ∼ β1°β2°β3°) as well as RUVBL2 and MscS (β1_GD_^S^β2^S^ and β4_GD_°β5°) are related via cross-fold sequence similarities. Additionally, we demonstrate transitive homology relationships: 1) between the OB fold of uL2 and the OB fold of RUVBL2 (Supplementary File 1, Table S1); and 2) between the SH3 fold of aS4 and the SH3 fold of MscS (Supplementary File 1, Table S2).

Thus, accounting for both cross-fold sequence similarity and transitive homology data, we can describe the general relationship between SH3 and OB folds as follows: 1) β1°β2°β3° is equivalent to β2^S^β3^S^β4^S^; 2) β4_GD_°β5° is equivalent to β1_GD_^S^β2^S^; 3) the GD-box motifs in β4_GD_° and in β1_GD_^S^ are homologous; 4) a duplicated SH3 **β1**_**GD**_^**S**^**β2**^**S**^**β3**^**S**^**β4**^**S**^β5^S^**-β1**_**GD**_^**S**^**β2**^**S**^β3^S^β4^S^β5^S^, after adaptation results in the emergence of an OB fold. **β2**^**S**^**β3**^**S**^**β4**^**S**^**β1**_**GD**_^**S**^**β2**^**S**^, renumbered N to C, gives β1°β2°β3°β4_GD_°β5° (Figure 4d).

We previously observed that SH3 and OB folds share two conserved supersecondary structural motifs: β1°β2°β3° ∼ β2^S^β3^S^β4^S^ and β4_GD_° ∼ β1_GD_^S^ (34). These conserved motifs are circularly permuted in SH3 with respect to OB. Here, we identify cross-fold sequence similarity between β5° and β2^S^, which are not structurally conserved. Structural variability of β5° as well as differences in structure between β5° and β2^S^ can be attributed to refolding, as part of the creative destruction process (see Discussion).

### OB and CLB Folds

The CLB fold, with six β-strands, is larger and more complex than the OB fold; the number and linkages of β strands differ between these folds. However, commonalities between OB and CLB folds with Alanine Racemase C topology in ECOD suggest ancestry via creative destruction. Several distinct regions of some OB and CLB fold proteins show cross-fold sequence similarities and partially superimpose in three dimensions (Figure 5e and 5f). These folds share one region of cross-fold sequence similarity corresponding to β1°β2°β4_GD_° and β1^C^β2^C^β3_GD_^C^, and a second region of cross-fold sequence similarity corresponding to β1°β2°β3°β4_GD_°L° and β4^C^L^C^β5^C^β6_GD_^C^α7^C^.

**Figure 5.**
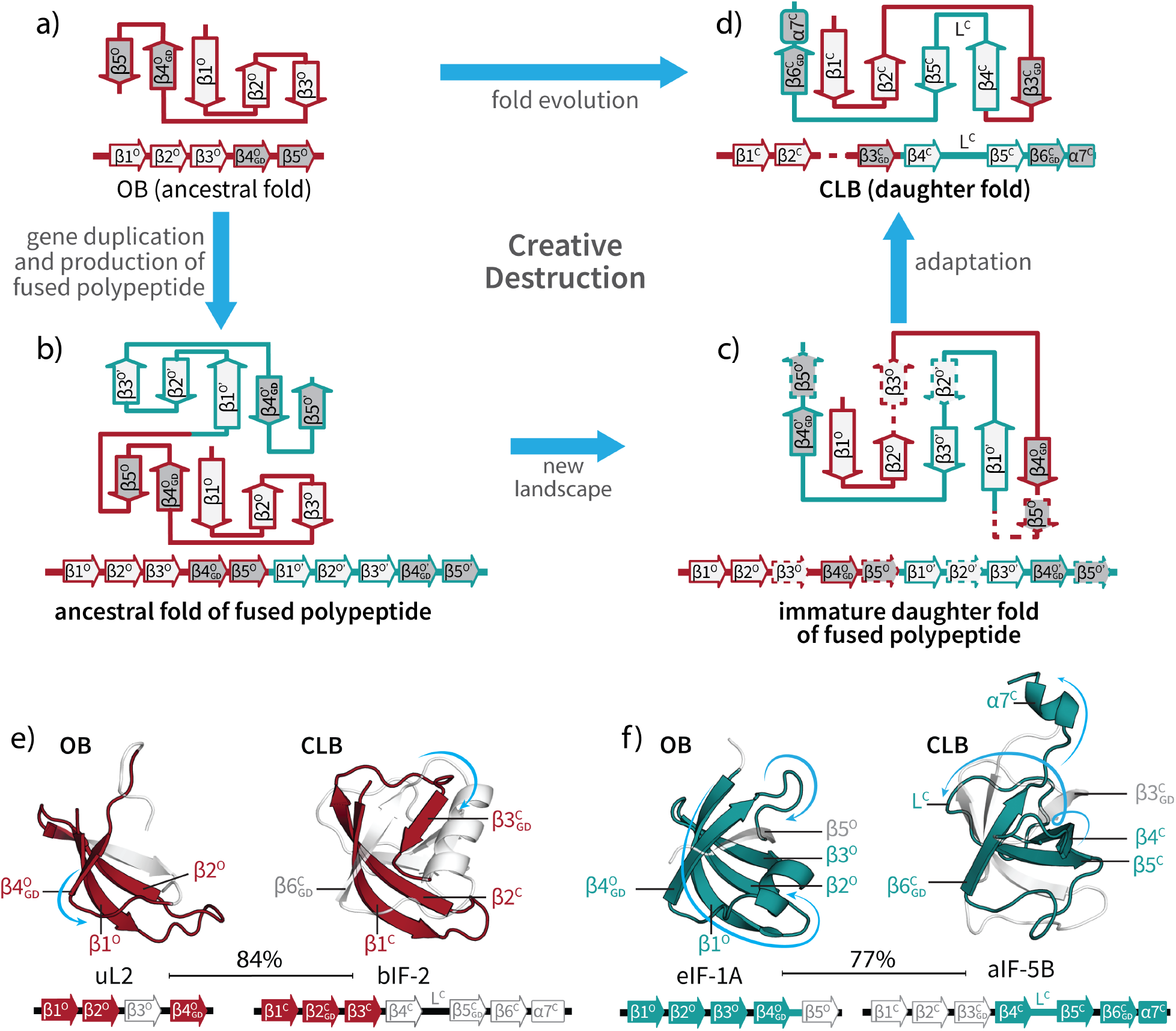
Conversion from OB Fold to CLB Fold by creative destruction. This figure shows, on a topological level, (a) the ancestral fold of the native protein, (b) the ancestral folds of the fused polypeptide, (c) the immature daughter fold in the fused polypeptide, and (d) mature daughter fold. A duplication of SH3 gives the fused polypeptide OB-OB. The fused polypeptide refolds and resolves by adaption, yielding a CLB fold. β4° ∼ β3^C^, and β4° ∼ β6^C^ are shaded dark gray to allow tracking of their positions during conversion from ancestral fold to daughter fold. (e) Cross-fold sequence similarity between uL2 and bIF2 corresponds to β1°β2°β4 ° and β1^C^β2^C^β3_GD_^C^. (f) Cross-fold sequence similarity between eIF-1A and aIF-5B corresponds to β1°β2°β3°β4_GD_ °L° and β4^C^L^C^β5^C^β6_GD_^C^α7^C^. Mapping of sequence similarity between OB and CLB: A bar between two folds indicates cross-fold sequence similarity, as given by HHalign scores. Regions that yield no sequence similarity are shown in white. Secondary structural elements that share sequence similarity are indicated by the same color in both members of the pair. Curved arrows follow secondary structural elements that are display differences in conformation between folds.

Shared features between OB and CLB folds are predicted by creative destruction. A duplicated and refolded OB fold, after adaptation, can give a CLB fold (Figure 5a-d). A duplication of β1°β2°β3°β4_GD_°β5° gives the fused polypeptide **β1**^**O**^**β2**^**O**^β3°**β4**_GD_^**O**^β5° -- **β1**^**O**^’**β2**^**O**^**’β3**^**O**^**’β4**_**GD**_^**O**^’β5°’ (where the prime indicates copy 2). This conversion to CLB retains the bold secondary elements, which are renumbered β1^C^β2^C^β3_GD_^C^β4^C^L^C^β5^C^β6_GD_°α7^C^ in the final product. The fused polypeptide refolds by extruding β5° and β5°’. β3° is lost and β2°’ refolds into a loop. β4°’ forms the terminus of the β-sheet, next to β1°. This model predicts cross-fold sequence similarities of β1^C^ ∼ β1°, β2^C^ ∼ β2°, β3_GD_^C^ ∼ β4_GD_°, β4^C^ ∼ β1°, L^C^ ∼ β2°, β5^C^ ∼ β3°, β6_GD_^C^ ∼ β4_GD_°, α7^C^ ∼ L°.

### Similarity between uL2 and bIF2

Cross-fold similarities are observed between the OB fold of uL2 and the CLB fold of bacterial Initiation factor 2 (bIF2). A region of similarity (HHalign score 84%) corresponds to β1°β2°β4_GD_° ∼ β1^C^β2^C^β3_GD_^C^. β4_GD_° ∼ β3_GD_^C^ have a homologous GD-box motif. In structure, a β- hairpin formed by β1°β2° is similar to the β-hairpin formed by β1^S^β2^S^. Strand β3^S^ has no equivalent in CLB.

### Similarity between eIF-1A and aIF5B

Cross-fold similarity is detected between the entire OB fold of initiation factor eIF-1A and the CLB fold in aIF-5B. Cross-fold sequence similarities (HHalign score 77%) between OB and CLB are: β1° ∼ β4^C^, β2° ∼ L^C^; β3° ∼ β5^C^; β4 °∼ β6 ^C^; β3°β4 ° ∼ β5^C^β6 ^C^; and L° ∼ α7^C^. A GD-box sequence motif is located in the loops of β4 ° ∼ β6_GD_^C^.

### Ancestral relationship between OB and CLB folds

The CLB fold comprises a large group of folds that include six-stranded β-barrels with a variety of topologies (37, 38). Here we analyzed CLB folds of the Alanine Racemase C topology. The cross-fold sequence similarity of the OB fold can be mapped to two distinct regions of this CLB fold (Figures 5e, 5f). Compelling homology relationships within OB folds (HHscore of 52-99%, Supplementary File 1, Table S1) and within CLB folds (HHscore of 84%, Supplementary File 1, Table S3) suggest that cross-fold similarity between OB and CLB folds can be generalized as follows: 1) β1^C^β2^C^β3_GD_^C^ ∼ β1°β2°β4_GD_°; 2) β4^C^L^C^β5^C^β6_GD_^C^α7^C^ ∼ β1°β2°β3°β4_GD_°L°; and 3) a duplicated four-stranded OB **β1**^**O**^β2°**β3**^**O**^**β4**_**GD**_^**O**^β5°--**β1**^**O**^**β2**^**O**^**β3**^**O**^**β4**_**GD**_^**O**^**L**^**O**^β5° results in the CLB topology β1^C^β2^C^β_GD_3^C^β4^C^L^C^β5^C^β6_GD_^C^α7^C^.

## Discussion

Protein structure is generally more conserved over evolution than sequence (26). The antipode; sequence conservation between divergent folds (cross-fold sequence similarities) suggests precipitous fold evolution (16), outpacing changes in sequence. We identified cross-fold sequence similarities between three β-barrel folds: SH3, OB, and CLB. Here we provide a basis of cross-fold sequence conservation in the form of a mechanism of fold evolution that interconverts SH3, OB and CLB folds.

Patterns of cross-fold sequence similarities between SH3 and OB proteins suggest that the OB fold arose via duplication of an SH3 gene, followed by production and folding of the fusion protein in a new landscape, then adaptation (34). Two SH3 genes fused and collapsed to a nascent OB fold that excluded terminal secondary elements. The mature OB fold was accomplished by loss of the terminal elements along with tuning mutations. OB fold proteins retain sequence and structure imprints from ancestral SH3 fold proteins. The polarity (SH3 -> OB versus OB -> SH3) is not fully resolved here and does not affect the conclusion. However, it seems most probable that SH3 is the ancestor of OB.

Patterns of cross-fold sequence similarity, identified here, suggest the CLB fold arose from OB gene duplication, folding of the fusion protein in a new landscape, and adaption. The final maturation of the CLB fold required loss of internal elements and tuning mutations. CLB fold proteins retain both sequence and structural imprints of OB fold proteins.

### Creative destruction

The combined data support a model of creative destruction in which ancestral folds readily beget daughter folds. Each step of creative destruction is known to be independently accessible and relatively frequent. An initial gene fusion step can cause insertion of one coding sequence in tandem with another (20, 39). It is assumed for simplicity that both ancestral genes express stable protein domains. A second step is expression of the fused polypeptide. The fusion alters the sequence and spatial context of the ancestral polypeptides and destabilizes the ancestral domains. At the same time alternative folding landscapes are opened. The fused polypeptide collapses to a daughter fold that differs from either of the ancestral folds.

### Motif Inheritance

Fold evolution by creative destruction is a funnel, not a random search of fold space. Daughter folds are derived in partially conservative and easily accessible processes from ancestral folds. Daughter folds are contingent on ancestral folds and inherit sequence and structural elements from them.

The GD-box demonstrates surprising persistence between SH3, OB and CLB folds. The presence of these motifs appears to provide stabilization of the β-barrel folds by bringing together a β-turn and a non-contiguous β-strand (33). SH3 and OB have one GD-box motif, and CLB has two (Figure 2). The creative destruction model explains the locations of these motifs in the sequence. Creative destruction does not require complex and improbable shuffling processes or transfer of isolated folding-incompetent motifs between proteins, as suggested by some previous models of fold evolution (28, 40, 41).

### Fold Plasticity

In creative destruction, a gene that encodes a protein with secondary elements ABCD can fuse with a gene that encodes a protein with secondary elements EFGH. The result is a fused polypeptide with ancestral secondary structure A**BC**D – E**FG**H. The fused polypeptide collapses and resolves by adaptation to daughter fold BCFG, which refolds to BCKG. The topology has been rewired, and F has changed conformation to K. Ancestral elements A, D, E, and H are lost. The most frequent conformational changes are expected to be conversion of α- helices to β-strands and vice versa (16). Some secondary elements of ancestors are retained while others are modified or lost. Conformational plasticity enables the integration of folding-competent sequences into new supersecondary structures.

Creative destruction is seen here to explain patterns of conserved and non-conserved sequence and structure between OB, SH3 and CLB folds. Our data suggest conformational changes: β2° converts to a loop in CLB (as exemplified by cross-fold sequence similarity between uL2 and bIF2, figure 5e), and a loop of the second copy of OB transforms to α7^C^ (as demonstrated by cross-fold sequence similarity between eIF-1A and aIF5B, Figure 5f). We also note an absence of β5 in the daughter CLB fold compared to the ancestral OB fold. Similarly, β5 in the ancestral SH3 fold is absent from the daughter OB fold. It is possible that the ancestral folds were four-stranded. A second possibility is that the ancestors were five-stranded, but β5 was destroyed during the creative destruction process, which could have occurred either during the duplication and fusion event or at a later step during adaptation. The model provides a mechanism of conversion from one protein fold to another, and offers a basis for establishing evolutionary relationships among diverse folds.

### Circular Permutation

Circular permutation has been documented previously in hundreds of proteins including concanavalin A (42), and other lectins (39), saposins (43), DNA methyltransferases (22, 44), and zinc ribbons (21, 45, 46) Two proteins related by circular permutation differ by two connections between secondary elements, but are otherwise fully conserved. Circular permutation can arise by two distinct mechanisms. One mechanism, seen only for concanavalin A, occurs at the polypeptide level (42) and is a post-translational re-wiring of the connections between secondary structural elements: 1) the N- and C-termini are joined; and 2) and a loop is cleaved, generating new N- and C-termini. The more general mechanism occurs on the gene level (22, 39), and as proposed here, involves creative destruction. Although the results appear indistinguishable, the two mechanisms are fundamentally different.

Circular permutation by creative destruction results from gene duplication, protein expression, exploration of new folding landscapes, and adaptation (Figure 1). The fused polypeptide product collapses to a new fold and adapts by loss of terminal elements. In circular permutation, the new folding landscape arrives at secondary elements inherited from the ancestral domains albeit with different linkages. Even though the ancestral domains and the daughter domain of circularly permuted proteins have common secondary structural elements, secondary elements of each ancestral domain are destroyed during the process.

We previously noted, for example, that archaeal and bacterial and versions of universal ribosomal protein uL33 are related by circular permutation (Figure 1e-h) (21, 46, 47). uL33 is a zinc ribbon, with two amphipathic β-hairpins (antiparallel ββ) linked by a zinc ion. The ordering of the ββ elements is switched in archaeal uL33 compared to bacterial uL33 (β1^a^β2^a^ ∼ β3^b^β4^b^ and β3^a^β4^a^ ∼ β1^b^β2^b^). Creative destruction provides a simple mechanism of conversion of archaeal to bacterial uL33 (and vice versa). Duplication of β1^a^β2^a^β3^a^β4^a^ gives fused polypeptide β1^a^β2^a^**β3**^**a**^**β4**^**a**^ -- **β1**^**a**^**β2**^**a**^β3^a^β4^a^. The fused protein collapses to a new fold, which omits ancestral elements β1^a^β2^a^ and β3^a^β4^a^, retaining the bold secondary elements to give daughter β1^b^β2^b^β3^b^β4^b^, where β1^a^ ∼ β3^b^, β2^a^ ∼ β4^b^, β3^a^ ∼ β1^b^ and β4^a^ ∼ β2^b^. The daughter fold adapts by loss of ancestral sequences for β1^a^β2^a^ and β3^a^β4^a^.

### Fold evolution by creative destruction

In previous models of fold evolution, cross-fold sequence similarities have been attributed to hypothetical ancestral peptides that are smaller than domains and are ancestral to them (11, 40, 48). Under these models, fold evolution depends on assembly, cooption, (41, 49) repetition (50) and decoration of small elements outside the context of a domain (11, 28, 40, 48) to generate new folds. Creative destruction by contrast acts on the level of domains and depends on fold plasticity, which is the facile exploration of new folding landscapes (51), and on a tendency of essentially any polypeptide sequence to self-associate. Robust folding is seen experimentally in the OB domain of uL2 and the CLB domain of bIF2 (52). Creative destruction resolves cross-fold similarities by a biologically plausible mechanism and suggests that the universe of protein folds is better described as a network than as a tree (53).

## Methods

### Representative SH3, OB and CLB domains of proteins from the translation system

Here, we studied the evolutionary relationship between the most prevalent and simplest β-barrel folds observed within the translation machinery: OB, SH3 and CLB. The structural information for specific representative proteins (uL2, aS4, and aS28, bIF2, aIF-5B) that contain these folds is summarized in Table 1. MSAs of these representative proteins containing orthologous proteins from Sparse and Efficient Representation of Extant biology (Bernier 2018 MBE) were obtained from ProteoVision (54). The MSAs of ribosomal proteins were trimmed to the domain boundaries as defined by the Evolutionary Classification of Domains, ECOD (25) and by Phase of Ribosomal evolution (1). The MSAs for ribosomal protein domains (Supplementary Files 2-6) were transformed to profile Hidden Markov Models using the HHsuite version 3.3.0 (55).

**Table 1.** Representative SH3, OB and CLB folds.

| **Table 1. Representative SH3, OB and CLB folds** |  |  |  |
| --- | --- | --- | --- |
| **Protein name** | **Fold Name** | **PDB code and chain** | **PDB fold range** |
| uL2 | OB | 1VY4, chain BA | 74-119 |
| aS4 | SH3 | 4V6U, chain AE | 180-243 |
| aS28 | OB | 4V6U, chain M | 2-71 |
| bIF-2 | CLB | 1ZO1, chain B | 190-267 |
| aIF-5B | CLB | 4V8Z, chain EC | 477-559 |

### Finding transitive relationships and cross-fold sequence similarity

To understand the relationships of SH3, OB, and CLB within and across folds, we searched the ECOD database (25). Profile files were retrieved from ECOD v283 (http://prodata.swmed.edu/ecod/distributions/ecod.develop283.hhm_db). An initial sequence similarity search to identify sequence similarity was performed using HHsearch with our SH3, OB, and CLB profiles as queries. To distinguish transitive homology relationships from cross-fold sequence similarities according to the hierarchical classification of ECOD the following criteria were applied: 1) HHalign scores greater than 50% within the same X-, H- and T-level groups in ECOD were considered transitive homologous relationships; 2) Protein domains yielding

### HHalign scores greater than 60% within different X-, H- or T-level groups in ECOD were considered cross-fold sequence similarities

Profiles of domains displaying either transitive homologous relations (Supplementary File 1, Table S1-S3) or cross-fold similarities (Supplementary File 1, Table S4 and S5) were retrieved and were compared in an all-versus-all fashion using HHalign with default parameters (55). HHalign scores of pairwise comparisons were deposited in a 2×2 matrix (Supplementary File 1, Figure S1).

### Structural analysis

Topology diagrams for these coordinate files were generated in ProteoVision using PDBsum (56). Three-dimensional representations of specific folds in the selected PDB files (Table 1) were rendered in PyMol (57). Pairs of folds displaying cross-fold sequence similarity were superimposed using Click (58) and manually adjusted using the pair fitting option in PyMol. Residues involved in cross-fold sequence similarities were highlighted by various colors in the topology diagrams and three-dimensional structure representations.

## Data Availability

Sequence alignments associated with this manuscript have been deposited in the FigShare repository 10.6084/m9.figshare.19412180.

## Acknowledgements

The authors thank Jessica Bowman, Petar Penev and Eric Smith for insightful discussions. CA-C was supported by the NASA Postdoctoral Program. This work was funded by the National Aeronautics and Space Administration grant 80NSSC18K1139 awarded to LDW and ASP.

