## Supplementary File 1 for "The Evolution of Protein Folds by Creative Destruction"

### Supplementary Methods

Multiple sequence alignments (MSAs) of proteins of the translation system were calculated using species from a Sparse and Efficient Representation of Extant biology (Bernier, et al. 2018). The MSAs of uL2, aS4, aS28, bIF-2 and aIF-5B were trimmed to the domain boundaries as defined by the Evolutionary Classification of Domains. Additionally, uL2, aS4 aS28 were trimmed to Phase of Ribosomal evolution (Kovacs, et al. 2017). Profile Hidden Markov Models (profiles) were built from trimmed MSAs of the domains with HHmake (Steinegger, et al. 2019) filtering columns with more than 50% of gaps. Profiles in HHM file format were retrieved from ECOD (Cheng, et al. 2014) version 283. Pairs of profiles were compared using HHalign (Steinegger, et al. 2019) with default parameters.

#### Figure S1. Cross-fold sequence similarity between SH3, OB and CLB folds

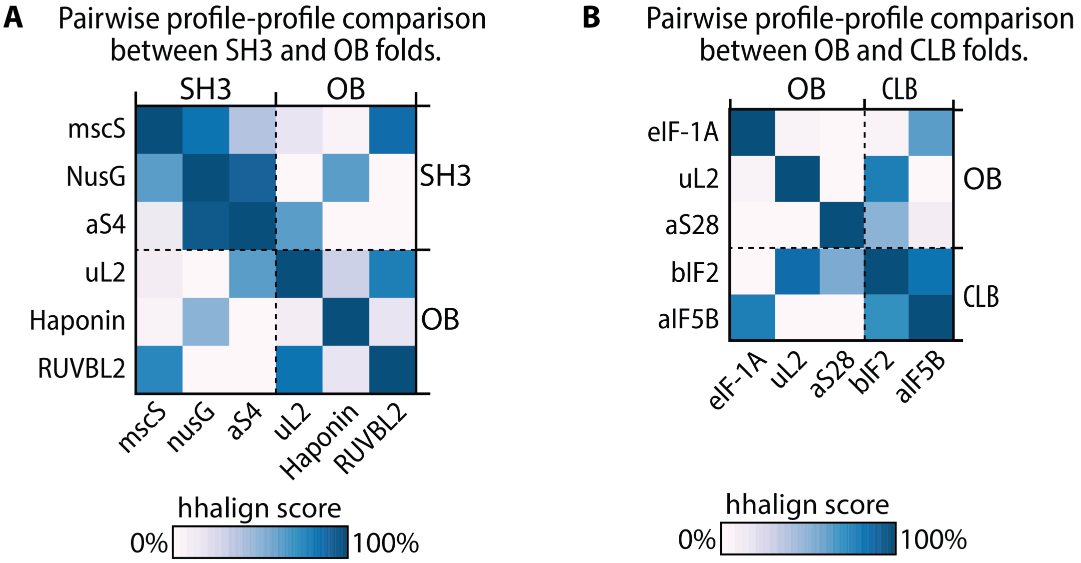


**Evolutionary relationship between beta-barrel folds.** HHalign scores of pairwise profile-profile comparisons represented by a color gradient in a two-by-two matrix. (A) Pairwise comparisons between SH3 and OB folds. (B) Pairwise comparisons between OB and CLB folds.

#### Table S1. Transitive homology relationships between OB domains

Transitive relationships between aS28 and uL2. Pairwise relationships in this table include: aS28, e1ny4A1, e2dgyA1, e3fp9A2, e2cqaA1, and uL2.

| Pair | Query_OB | Target_OB | hhalign_score | Aligned_cols |
|---|---|---|---|---|
| 1 | aS28 | e1ny4A1 | 99.94 | 70 |

```
Pair 1.
>e1ny4A1 A:1-71 000000346
Probab=99.93  E-value=4.3e-36  Score=171.28  Aligned_cols=70  Identities=99%  Similarity=1.412  Sum_probs=67.0
Template_Neff=3.300

Q aS28              2 AEDEGYPAEVIEI--IGRTGTTGDVTQVKVRILEGRDKGRVIRRNVRGPVRIGDILILRETERE-AREIKSRR   71 (71)
Q Consensus         2 ~~~~~~pAeViei~~vGrTG~~GEv~qV~~rileG~dkGrii~RNv~GPvr~gDIl~LrETerE~A~~i~~rr   74 (74)
                      .+++.++|+|++|  +||||++|+++||+|++|+|.|+||++.|||+||||+||||+|.||||| |++|..++
T Consensus         2 ~e~~~~~A~V~kV--lGRTGs~G~~tQVrve~l~~~~~~R~iiRNVKGPVr~GDil~LlEsERE-Arrl~~~~   71 (71)
T e1ny4A1           2 AEDEGYPAEVIEI--IGRTGTTGDVTQVKVRILEGRDKGRVIRRNVRGPVRVGDILILRETERE-AREIKSRR   71 (71)
T ss_pred            CCCCCCceeEEEE--eecccccceeEEEEEEeecCCCceeeeeccCcccccceeeeeehhHHH-HhhhhccC
```

```
Confidence           4567899999999  99999999999999999999999999999999999999999999999999 99998775
```

| | | | | |
|---|---|---|---|---|
| 2 | e1ny4A1 | e2dgyA1 | 52.33 | 48 |

```
Pair 2.
>e2dgyA1 A:1-111 000161818
Probab=52.33  E-value=0.00087  Score=22.05  Aligned_cols=48  Identities=19%  Similarity=0.241  Sum_probs=26.0
Template_Neff=6.100

Q ss_pred              CCCCCceeEEEEeeccccccceeEEEEEEeecCCCceeeeeecc------CcccccceeeeeehhH
Q e1ny4A1            3 EDEGYPAEVIEIIGRTGTTGDVTQVKVRILEGRDKGRVIRRNVR------GPVRVGDILILRETE   61 (71)
Q Consensus          3 e~~~~~A~V~kVlGRTGs~G~~tQVrve~l~~~~~~~R~iiRNVK------GPVr~GDil~LlEsE   61 (71)
                       ++....+.|++.+|..         ..+|+.-+    +..++..+.       =+|.+||.+.+.-++
T Consensus        13 ~e~e~~G~V~~~lGn~----------V~~~d----G~~~l~~i~GK~Rk~i~I~~GD~V~ve~~~   66 (111)
T e2dgyA1          13 SNQQQIVRVLRTPGNN-------LHEVETAQ----GQRFLVSMPSKYRKNIWIKRGDFLIVDPIE   66 (111)
T ss_pred              CCCceeEEEeecCCCc-------eeEEEecc----ceeeeeccccccceeeeecCceeEeeecc
Confidence             3445568888887742       12232222    122233333      348999998776554
```

| | | | | |
|---|---|---|---|---|
| 3 | e2dgyA1 | e3fp9A2 | 95.48 | 63 |

```
Pair 3.
>e2dgyA1 A:1-111 000161818
Probab=91.29  E-value=2.1e-06  Score=36.02  Aligned_cols=63  Identities=16%  Similarity=0.080  Sum_probs=47.7
Template_Neff=6.100

Q ss_pred              hhhhhhhhcCCceeEeeccchhHHHHhhhcccccCCCCCCcccccCCCCCCCCCCceeeeecc------cceeeeeccc
Q e3fp9A2            7 STLREILADGHRALVVGHADEERVVWLADPLIAEDLPDGLPEALNDDTRPRKLRPGDSLLVDTK------AGYAFERIPK  80 (92)
Q Consensus          7 vtv~Evl~d~~R~lV~~~~deervv~lA~~L~~~~~~~~~~~~~~~~~~~~~~~~Lr~GDsllvD~r------sg~A~E~Vpk  80 (92)
                       .+|.+.++. .++.|-...+..+++++.+.+...                    ..+.+||.|+|++.      .|....+.++
T Consensus        19 G~V~~~lGn-~~~~V~~~dG~~~l~~i~GK~Rk~----------------i~I~~GD~V~ve~~~~~~~~~kg~I~~~~~r  81 (111)
T e2dgyA1          19 VRVLRTPGN-NLHEVETAQGQRFLVSMPSKYRKN----------------IWIKRGDFLIVDPIEEGEKVKAEISFVLCK  81 (111)
T ss_pred              EEEeecCCC-ceeEEEeccceeeeeecccccce----------------eeeecCceeEeeeccccccceeeeeeech
Confidence             567888875 688998889999999999998651                4589999999874      3556666666


Q ss_pred              hhhhhh
Q e3fp9A2           81 AEVEDL   86 (92)
Q Consensus         81 ~eve~L   86 (92)
                       .++..|
T Consensus        82 ~~v~~L   87 (111)
T e2dgyA1          82 DHVRSL   87 (111)
T ss_pred              hHHHHH
Confidence             655543
```

| | | | | |
|---|---|---|---|---|
| 4 | e3fp9A2 | e2cqaA1 | 75.73 | 42 |

```
Pair 4.
>e3fp9A2 A:60-151 000136897
Probab=75.73  E-value=2.4e-05  Score=30.04  Aligned_cols=42  Identities=21%  Similarity=0.359  Sum_probs=34.5
Template_Neff=3.900

Q ss_pred              eeeeecccceeeechhhHHHH-------------HHhhhhcccCceEEEEeccccee
Q e2cqaA1           30 LTLKTTEMETIYDLGTKMIE-------------SLTKDKVQAGDVITIDKATGKI   71 (82)
Q Consensus         30 itLkT~d~e~~l~lg~~i~e-------------~l~kekV~~GDVI~Id~~sG~V   71 (82)
                       |..-..+++....|...+..              .+.++++++||-+.+|+.+|-.
T Consensus        20 lV~~~~deervv~lA~~L~~~~~~~~~~~~~~~~~~~~~Lr~GDsllvD~rsg~A   74 (92)
T e3fp9A2          20 LVVGHADEERVVWLADPLIAEDLPDGLPEALNDDTRPRKLRPGDSLLVDTKAGYA   74 (92)
T ss_pred              eEeecccchhHHHHhhhcccccCCCCCCcccccCCCCCCCCCCceeeeeeccccee
Confidence             666778889999998888875              34558999999999999854
```

| | | | | |
|---|---|---|---|---|
| 5 | e2cqaA1 | uL2 | 80.13 | 46 |

```
Pair 5.
>uL2
Probab=80.13  E-value=1.4e-05  Score=27.76  Aligned_cols=46  Identities=20%  Similarity=0.289  Sum_probs=31.9
Template_Neff=4.991

Q ss_pred              eeeeeeeEeeeecCCCCCCcceeeeeeeeec---cceeeechhhHHHHHHhhhhcccCceEEE
Q e2cqaA1            6 IIEGEVVEIQIDRPATGTGSKVGKLTLKTT---EMETIYDLGTKMIESLTKDKVQAGDVITI   64 (82)
Q Consensus          6 V~EGEV~ei~i~~p~~g~~~~~~~~itLkT~---d~e~~l~lg~~i~e~l~kekV~~GDVI~I   64 (82)
                       .+.|.|.+|.-|.--+.   ..  .-++-.   |+++.|-|.|+        |+.+||.|++
T Consensus         2 ~~~g~V~~I~hdP~R~A---pi--A~v~~e~~~dGe~~yilApe--------G~~vG~~i~~   50 (51)
T uL2               2 GIPAKVAAIEYDPNRSA---RI--ALLHYV---DGEKRYIIAPD--------GLQVGQQVVA   47 (48)
Confidence             36789999988862111   11  233444   88999988886        9999999874
```

#### Table S2. Transitive homology relationships between SH3 domains

Transitive relationships between e2oauA1 and aS4. Pairwise relationships in this table include: e2oauA1, e4agfA10, e1nz9A1, and aS4.

<table>
<tr><th>Pair</th><th>Query_SH3</th><th>Target_SH3</th><th>hhalign_score</th><th>Aligned_cols</th></tr>
<tr><td>1</td><td>e2oauA1</td><td>e4agfA10</td><td>100.0</td><td>67</td></tr>
<tr><td colspan="5"><pre>
Pair 1.
>e4agfA10 A:113-179 001172576
Probab=99.96  E-value=4.3e-34  Score=137.92  Aligned_cols=67  Identities=99%  Similarity=1.391  Sum_probs=65.8
Template_Neff=6.800

Q ss_pred               CccchhhhhhhhHhhcccccceeEEecCceeeEEEEeeeEEEEeecCCeeEEeecccceeeeeeeecC
Q e2oauA1            1 GSLSNLAAGVLLVMFRPFRAGEYVDLGGVAGTVLSVQIFSTTMRTADGKIIVIPNGKIIAGNIINFS   67 (67)
Q Consensus          1 d~l~n~~agi~i~~~~~~~~vGD~I~i~~~~~G~V~~i~l~~t~l~~~dg~~v~IPNs~l~~~~v~N~s   67 (67)
                       |+++|++||++|++.+||++||||+++|..|+|++|+|++|++++.||+.++|||+.+.++++.|||
T Consensus          1 d~l~n~~sGi~i~~~~pf~vGD~I~i~~~~~G~V~~I~l~~T~l~~~dg~~v~IPNs~l~~~~i~N~s   67 (67)
T e4agfA10           1 GSLSNLAAGVLCVMFRPFRAGEYVDLGGVAGTVLSVQIFSTTMRTADGKIIVIPNGKIIAGNIINFS   67 (67)
T ss_pred               CccchhhhhhhhhhhcccccceeEEecCceeeEEEEeeeEEEEeecCCeeEEeecccceeeeeeeecC
Confidence             6899999999999999999999999999999999999999999999999999999999999999997
</pre></td></tr>
<tr><td>2</td><td>e4agfA10</td><td>e1nz9A1</td><td>95.31</td><td>43</td></tr>
<tr><td colspan="5"><pre>
>e4agfA10 A:113-179 001172576
Probab=95.31  E-value=1.2e-06  Score=28.25  Aligned_cols=43  Identities=23%  Similarity=0.312  Sum_probs=28.7
Template_Neff=6.800

Q ss_pred               cccccCceEEEeeccCCCccceeeEeecCCCceeEEEEeeccccceeeeecccce
Q e1nz9A1            3 VAFREGDQVRVVSGPFADFTGTVTEINPERGKVKVMVTIFGRETPVELDFSQV   55 (58)
Q Consensus          3 ~~~~~Gd~V~I~~Gpf~g~~g~v~~v~~~k~~v~V~v~~~gr~~~v~l~~~~v   55 (58)
                       -+|++||.|++-     |.+|.|.++....   +..-...|+  .+.++.+.+
T Consensus         16 ~pf~vGD~I~i~-----~~~G~V~~I~l~~---T~l~~~dg~--~v~IPNs~l   58 (67)
T e4agfA10          16 RPFRAGEYVDLG-----GVAGTVLSVQIFS---TTMRTADGK--IIVIPNGKI   58 (67)
T ss_pred               cccccceeEEec-----CceeeEEEEeeeE---EEEeecCCe--eEEeeccce
Confidence             468999999984     6899999998643   222344444  355555544
</pre></td></tr>
<tr><td>3</td><td>e1nz9A1</td><td>aS4</td><td>88.97</td><td>27</td></tr>
<tr><td colspan="5"><pre>
>e1nz9A1 A:1-58 000026343
Probab=90.78  E-value=1.2e-06  Score=31.32  Aligned_cols=27  Identities=30%  Similarity=0.420  Sum_probs=25.2
Template_Neff=7.100

Q SH3-1_eS04c_18     1 PFEKGAYVFVTQGKNVARKGRIVEIKR   27 (59)
Q Consensus          1 pfe~G~~v~I~~G~h~G~iGkI~~I~~   27 (73)
                       +|++|+.|.|++|.-.|..|+|.++..
T Consensus          4 ~~~~Gd~V~I~~Gpf~g~~g~v~~v~~   30 (58)
T e1nz9A1            4 AFREGDQVRVVSGPFADFTGTVTEINP   30 (58)
T ss_pred               ccccCceEEEeeccCCCccceeeEeecC
Confidence             478999999999999999999999976
</pre></td></tr>
</table>

#### Table S3. Pairwise homology relationships between CLB domains

<table>
<tr><th>Pair</th><th>Query_ARC</th><th>Target_ARC</th><th>hhalign_score</th><th>Aligned_cols</th></tr>
<tr><td>1</td><td>bIF-2</td><td>aIF-5B</td><td>84.99</td><td>84</td></tr>
<tr><td colspan="5"><pre>
Pair 1
>1_1G7S_1|Chain A|TRANSLATION INITIATION FACTOR IF2/EIF5B|Methanothermobacter thermautotrophicus (145262)_460-
587
Probab=84.99  E-value=6.3e-06  Score=36.54  Aligned_cols=78  Identities=18%  Similarity=0.245  Sum_probs=59.1
Template_Neff=4.396

Q bIF-2             15 GRGPVATVLVR-EGTLHKGDIVLC--G---FEYGRVRAMRNEL-GQEVLEAGPS--IPVEILGL---SGVPAAGDEVTVV  82 (101)
Q Consensus         15 grGpVATvLVQ-~GTL~vGD~vva--G---~~~GRVRAm~dd~~G~~v~eAgPS--~PVeilGl---~~vP~AGD~f~Vv  83 (102)
                       ..-+|.-|.|. .|+|++|..++-  |    ..-|+|+.|.|+. -+.+++|.-.   -.+.|-|-    .--=..||.++|-
T Consensus         20 s~PAIVGVeVl~~G~ir~~~~li~~~G~~~k~vG~I~~Iqd~~-~~~v~~A~~G~eVAiSI~G~~tvGRqI~egD~LYvd  98 (139)
T aIF-5B            18 SKPAIGGVEVL-TGVIRQGYPLMNDDG---ETVGTVESMQDKG--ENLKSASRGQKVAMAIKDA-VYGKTIHEGDTLYVD  90 (128)
Confidence             34467778899 999999999997  6   6789999998875 4777777544  45556554   2234689999999


Q bIF-2             83 RDEKKAR-EVA   92 (101)
Q Consensus         84 ~~e~~AR-eia   93 (102)
                       -.|+.|+ .+-
T Consensus         99 ipe~~~k~~l~  109 (139)
T aIF-5B            91 IPENHYH-ILK  100 (128)
Confidence             9999999 443
</pre></td></tr>
</table>

#### Table S4. Cross-fold sequence similarity between OB domains and SH3 domains

| Similarity pairs between and | | | | |
|---|---|---|---|---|
| **Pair** | **OB** | **SH3** | **hhalign_score** | **Aligned_cols** |
| 1 | uL2 | aS4 | 64.15 | 32 |

```
Pair 1.
>SH3-1_eS04c_186497_PYRFU_178-236
Probab=64.15  E-value=6.5e-05  Score=22.35  Aligned_cols=32  Identities=13%  Similarity=0.135  Sum_probs=26.5
Template_Neff=6.041

Q uL2               4 PAKVAAIEYDPNRSARIALLHYVDGEKRYIIA   35 (48)
Q Consensus         4 ~g~V~~I~hdP~R~ApiA~v~~edGe~~yilA   35 (48)
                      .|+|.+|+.-+.|...++.++-+||+++-.+.
T Consensus        19 ~G~I~~I~~~~~~~~~~~~V~ie~~~ge~f~T~~   50 (59)
T aS4              19 KGRIVEIKRFPMGWPDVVTIEDEEGELFDTLK   50 (59)
Confidence            58999999999999999999888998654443
```

| Pair | OB | SH3 | hhalign_score | Aligned_cols |
|---|---|---|---|---|
| 2 | RUVBL2(e2cqaA1) | MscS(e2oauA1) | 85.40 | 25 |

```
Pair 2.
>e2cqaA1 A:8-89 000000382
Probab=85.40  E-value=5.8e-06  Score=22.89  Aligned_cols=25  Identities=12%  Similarity=0.228  Sum_probs=21.3
Template_Neff=4.100

Q ECOD_000025611   15 FRPFRAGEYVDLGGVAGTVLSVQIF   39 (67)
Q Consensus       15 ~~~~~~gd~i~~~~~~~g~v~~i~~~   39 (67)
                     ......||.|.++...|.|.+++.-
T Consensus       53 kekV~~GDVI~Id~~sG~V~klGRs   77 (82)
T e2cqaA1         53 KDKVQAGDVITIDKATGKISKLGRS   77 (82)
T ss_pred            hhhcccCceEEEeccccceeeeeecc
Confidence          3577899999999999999998754
```

#### Table S5. Cross-fold sequence similarity between OB domains and CLB domains

| Similarity pairs of ECOD domains with sequence similarity between and | | | | |
|---|---|---|---|---|
| **Pair** | **OB** | **ARC** | **hhalign_score** | **Aligned_cols** |
| 1 | uL2 | bIF-2 | 87.14 | 36 |

```
Pair 1.
>OB-1_uL02b_262724_THET2/1-276_73-120
Probab=87.14  E-value=3.9e-06  Score=31.11  Aligned_cols=36  Identities=25%  Similarity=0.343  Sum_probs=31.6
Template_Neff=5.204

Q 1_bIF2_511145_    3 ASGAVIESFLDKGRG-PVATVLVREGT---------LHKGDIVLCG   38 (101)
Q Consensus         3 A~G~VIEa~LDkgrG-pVATvLVQ~GT---------L~vGD~vvaG   38 (101)
                      -.|+|++-.-|.+|. |+|.|...||+         |.+||.|.+|
T Consensus         3 ~~g~V~~I~hdP~R~ApiA~v~~edGe~~yilApeGl~vG~~I~~G   48 (48)
T OB-1_uL02b_262    3 IPAKVAAIEYDPNRSARIALLHYVDGEKRYIIAPDGLQVGQQVVAG   48 (48)
Confidence            4688999999986 89999999998         7899998876
```

| Pair | OB | ARC | hhalign_score | Aligned_cols |
|---|---|---|---|---|
| 2 | aS28 | bIF-2 | 75.90 | 39 |

```
Pair 2.
>OB-1_eS28c_186497_PYRFU_1-71
Probab=75.90  E-value=2.3e-05  Score=27.65  Aligned_cols=39  Identities=21%  Similarity=0.341  Sum_probs=29.2
Template_Neff=3.035

Q bIF2_262724_TH    8 KEAGPGSAVQVLGFQELPH----------------------------AGDVVEWVPDLEAAKEIAEE   46 (63)
Q Consensus         8 ~eAgPS~PVeVlGl~~vP~----------------------------AGD~f~Vv~dE~~AReIa~~   46 (63)
                      ..+.|..-+||+|-+|+-.                            -||.+..-++|..||.|.-.
T Consensus         4 ~~~~pAeVieivGrTG~~GEv~QVkcrIleG~dkGRIitRNv~GPVR~GDIlmLrETerEAr~i~~r   70 (71)
T OB-1_eS28c_186    4 DEGYPAEVIEIIGRTGTTGDVTQVKVRILEGRDKGRVIRRNVRGPVRIGDILILRETEREAREIKSR   70 (71)
Confidence            4567777788877766543                            58888999999999888644
```

| Pair | OB | ARC | hhalign_score | Aligned_cols |
|---|---|---|---|---|
| 3 | eIF-1A(e2oqkA1) | aIF-5B | 77.25 | 75 |

```
Pair 3.
>ECOD_000163439_e2oqkA1 | 2.1.1.20 | 2OQK A:22-115 | A: beta barrels, X: OB-fold, H: NO_H_NAME, T: Nucleic acid-
```

```
binding proteins, F: eIF-1a | Protein: Putative translation initiation factor eIF-1A
Probab=77.25  E-value=2e-05  Score=29.37  Aligned_cols=75  Identities=20%  Similarity=0.236  Sum_probs=53.9
Template_Neff=9.628

Q 1_1G7S_1|Chain   42 DGETVGTVESMQDKGENLKSASRGQKVAMAIKDAVYG-KTIHEGDTLYVDIPENHYHILKEQLSGDLTDEELDLMDKI  118 (128)
Q Consensus        42 ~Gk~vG~I~~Iqd~~~~v~~A~~G~eVAiSI~g~tvG-RqI~egD~LYvdipe~~~k~l~~~~~d~L~~de~e~L~e~  118 (128)
                      +|..+|+|.+..-++...-....|.+.-+++.|..=+ .-|..||.+.+++.+..-  -..++...++.++...|++.
T Consensus        10 ~~~~~g~V~~~~g~~~~~V~~~~g~~~~~~~~~~k~r~~~~i~~gd~V~i~~~~~~~~--~~~~~i~~~~~~~~~~~~l~~~   85 (94)
T ECOD_000163439   10 EGQEYGQVQRMLGNGRLDAYCFDGQKRLCHIRGKMRKKVWVNPGDIVLVSLRDFQD--SKGDIILKYTPDEARALKSK   85 (94)
Confidence            5788888888777665555555788888888888322 478899999998877652  23456677888887777553
```
